# Bio-orthogonal Mid-Infrared Photothermal Imaging Unveils Altered Stearoyl-CoA desaturase (SCD) Activity in Ovarian Cancer Cells

**DOI:** 10.1101/2025.07.02.662881

**Authors:** Xinyan Teng, Tianyu Xia, Mingsheng Li, Jiaze Yin, Jianpeng Ao, Guangrui Ding, Dashan Dong, Chinmayee V. Prabhu Dessai, Yinu Wang, Ana Maria Isac, Qing Xia, Daniela Matei, Hongjian He, Ji-Xin Cheng

**Author notes:** **Corresponding authors** (J.X.C.), (H.H.), (D.M.).

## Abstract

Stearoyl-CoA desaturase (SCD)-catalyzed fatty acid desaturation plays a key role in cancer metabolism reprogramming. While existing approaches enable quantification of SCD expression, directly monitoring *in situ* SCD enzymatic activity alterations remains challenging. Here, we report a strategy to measure intracellular SCD activity alterations and SCD product distributions by mid-infrared photothermal (MIP) imaging. Fully deuterated palmitic acid (PA-d31) is repurposed here as a chemical probe to report the SCD activity via a new peak at 2246 cm^−1^, corresponding to the stretching of D-C=C-D bonds in the SCD product. The MIP intensity ratio between the product-associated signal and the total C-D signal defines the apparent SCD activity index. By visualization of SCD-catalyzed metabolic conversion, our approach reveals altered SCD activity in cisplatin-treated ovarian cancer cells, ovarian cancer stem cells and cells fed with oleic acid. Non-uniform distributions of the SCD product are also observed. These results establish bio-orthogonal MIP microscopy as a platform for studying intracellular enzymatic reactions in living systems.

## Introduction

Cellular lipid metabolism dysregulation is increasingly recognized as a hallmark of ovarian cancer metabolic reprogramming (*1–4*). Among all lipid metabolic enzymes, stearoyl-CoA desaturase (SCD) is a critical enzyme that catalyzes the Δ9 desaturation of saturated fatty acids (SFAs), introducing a carbon-carbon double bond (C=C) into the hydrocarbon chain to generate the corresponding monounsaturated fatty acids (MUFAs) from palmitic acid (PA) and stearic acid (SA) (*5*). This biochemical reaction plays a central role in maintaining lipid homeostasis and supporting rapid membrane biosynthesis and stress adaptation in proliferating cells (*6*). Reprogrammed SCD activity has emerged as a key metabolic regulator and therapeutic target in ovarian cancer (*7–10*). Hence, understanding SCD activity alterations could unveil the metabolic plasticity and identify potential targets for ovarian cancer diagnosis.

Although essential, directly measuring SCD activity is still challenging. Conventional expression assays such as Western blot (WB) and qPCR can quantify SCD expression at the protein or mRNA level (*11–14*). However, these static expression measurements fail to capture the actual SCD activity. While mass spectrometry-based methods (*12, 13*) enabled sensitive quantification of SCD products, assessing enzymatic activity with single-cell resolution remains difficult. Although fluorophore-labeled fatty acid (FA) probes have been widely used to visualize FA metabolic trafficking (*15*), these probes fail to report desaturase activity because fluorophores are insensitive to FA structural changes regardless of the labeling position. Fluorescence-based methods such as RNA fluorescence *in situ* hybridization (RNA-FISH) and immunofluorescence (IF) can precisely localize the SCD gene, yet it does not provide spatial information about SCD activity (*16*).

Recently developed mid-infrared photothermal (MIP) microscopy, a vibrational imaging modality that provides molecular stretching information and sub-cellular resolutions (*17, 18*), has been applied to diverse biochemical fields, including the analysis of enzyme activities (*19*), elucidation of carbohydrate metabolic pathways (*20*), investigation of protein structures (*21, 22*), and exploration of FA metabolism (*23–25*). MIP imaging combined with small vibrational tags has been applied to monitor fatty acid metabolism in biological systems (*23–25*). These studies demonstrate the feasibility of combining MIP with vibrational tags to investigate metabolic activities. However, current MIP-based metabolism studies were predominantly focused on tracing the uptake and accumulation of exogenous substrates, rather than dynamically monitoring the metabolic activity using a responsive vibrational probe.

Here, to measure intracellular SCD activity at single-cell resolution and map the intracellular distribution of the SCD products, we employed MIP microscopy by repurposing the well-established fully deuterated palmitic acid (PA-d31) as a substrate and reporter of SCD activity, revealing a previously unrecognized functional role for this classic FA uptake tracer. This strategy exploits the bond-selective imaging capability of MIP microscopy. The SCD action on PA-d31 generates an unsaturated D-C=C-D motif within the SFA chain, producing a new vibrational peak that reflects the SCD activity. Importantly, the C-D vibrational signal lies within the cell silent vibrational window, where the endogenous biomolecules produce minimal background signal (*26*), enabling a highly specific detection of SCD activity.

Using a counter-propagated laser-scanning mid-infrared photothermal (MIP) microscopy **(Fig. 1A)**, we monitored the intracellular SCD activity through the emergence of a new peak at 2246 cm^−1^ **(Fig. 1B** and **C)**, corresponding to the C-D stretch of D-C=C-D generated by SCD activity. To extract the peak intensity from the intracellular water background, we applied asymmetrically reweighted penalized least squares (arPLS) baseline correction to the hyperspectral MIP images (*27*). Using this method, we directly measured the SCD activity alterations **(Fig. 1D)**. We observed an elevated SCD activity in cisplatin treated ovarian cancer cells and in cancer stem cells (CSCs). Interestingly, supplementation of cells with unsaturated fatty acids suppressed the SCD activity. Spatially, we found that SCD products preferentially localize to the surface of lipid droplets (LDs) rather than distributing uniformly, and in CSCs these products further contribute to plasma membrane biosynthesis.

**Figure 1.**
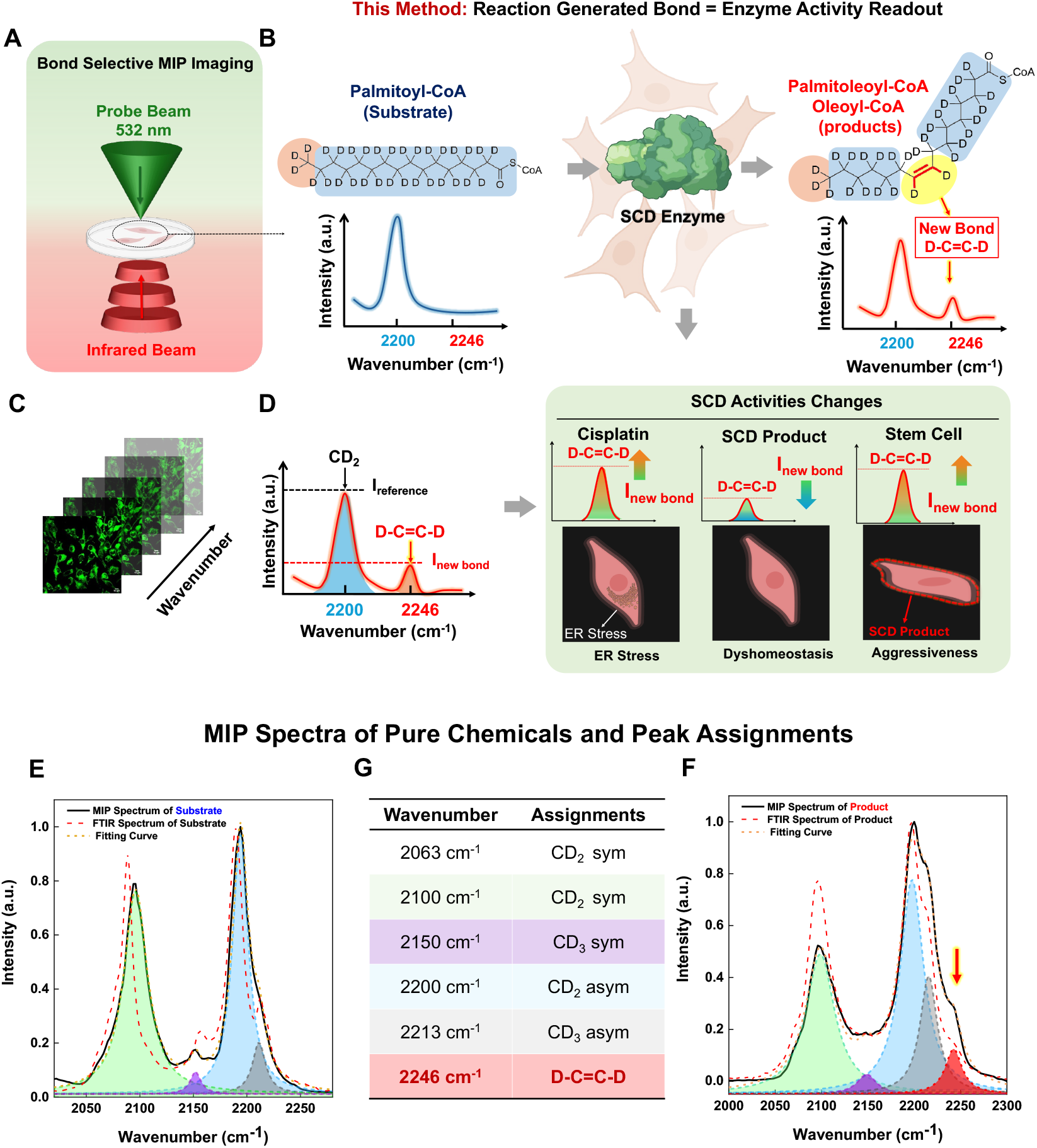
Measuring SCD Activity by D-C=C-D Generation through MIP Imaging. **(A)** Schematic of counter-propagation bond-selective MIP imaging of ovarian cancer cells. **(B)** Equation of SCD enzymatic reaction and scheme of vibrational signatures between SCD product and substrate. **(C)** Scheme of hyperspectral MIP imaging unveiling vibrational signatures of SCD activity. **(D)** Scheme summarizing SCD activity alterations under different biological conditions. **(E-G)** MIP spectra (black) and FTIR (dashed red) of pure PA-d31 **(E)** and OA-d34 **(F)** and their peak deconvolutions and peak assignments **(G)**. **(A)-(D)** were created in BioRender. He, H. (2026) https://BioRender.com/t1ssolv

## Results

### MIP Spectroscopy of Fatty Acid Desaturation

Stearoyl-CoA desaturase (SCD) is a pivotal enzyme responsible for catalyzing the conversion of saturated fatty acids (SFAs) into monounsaturated fatty acids (MUFAs), a process essential for maintaining the membrane fluidity and supporting the lipid synthesis (*5, 28*). The SCD activity introduces a double bond (C=C) within the SFA chain at the Δ9 position, resulting in a distinct vibrational signature associated with C=C bond formation.

We harnessed a laser-scanning MIP microscope to measure such vibrational signature. The schematic of MIP microscopy is shown in **Fig. 1A**. Briefly, a pump-probe strategy is used to detect the photothermal effects induced by infrared (IR) absorption (*17, 29, 30*). In this work, a 532 nm visible beam was used as the probe beam, while a tunable pulsed quantum cascade laser (QCL) served as the IR pump source. The pulsed IR laser excites specific molecular vibrations, causing a transient local temperature rise, followed by thermal expansion of particles and/or refractive index change. These photothermal effects are detected by the 532 nm focused visible probe beam. For particles with a diameter of 1.0 μm, the thermal diffusion length is about 200 nm (*30*). Thus, for lipids inside cells, MIP microscopy provides the IR absorption contrast at the spatial resolution of the visible beam.

Since palmitic acid (PA) is a key substrate of SCD and a primary source of FAs in cells (*1, 7*), fully deuterated palmitic acid (PA-d31) was used as an SCD activity substrate and repurposed as a SCD activity reporter. PA-d31 is first converted to saturated fatty acyl-CoA through acyl-CoA formation. SCD then catalyzes Δ9 desaturation of the saturated fatty acyl-CoA substrate to generate the corresponding monounsaturated fatty acyl-CoA product containing a Δ9 C=C double bond **(Fig. 1B)**. Using hyperspectral MIP imaging **(Fig. 1C)**, this D-C=C-D stretching mode provides a specific vibrational signature for reporting SCD activity under different biological conditions **(Fig. 1D)**.

We first examined the vibrational signature of SCD product by comparing MIP and Fourier transform infrared (FTIR) spectra of PA-d31 (SCD substrate) and OA-d34 (the Δ9-MUFA) (**Fig. 1E** and **1F**). Both MIP (solid curve) and FTIR (dashed red curve) spectra, following deconvolution (round dot curve), revealed vibrational signatures that distinguish the product from substrate. The most prominent feature is the emergence of a new peak at 2246 cm^−1^ in the product spectra.

To evaluate the detection sensitivity of our MIP microscope, we then tested the limit of detection (LOD) of C-D vibration at 2200 cm^−1^, and the minimum detectable product fraction which is defined as the percentage of OA-d34 in a mixture of OA-d34 and PA-d31 based on the peak at 2246 cm^−1^. Results show that the LOD of the C-D signal is 3.31 μM OA-d34 (**Fig. S1**). The MIP peak intensity at 2246 cm^−1^ is linear to the product fraction (**Fig. S2**). The minimum detectable product fraction was estimated to be 6% (molar ratio) OA-d34 in the substrate background (**Fig. S2**). These results suggest that the distinctive new peak at 2246 cm^−1^ enables precise and sensitive monitoring of the SCD activity.

### MIP imaging of intracellular SCD enzymatic activity

Next, we applied this strategy to measure intracellular SCD activity. We performed hyperspectral MIP imaging from 2000 cm^−1^ to 2300 cm^−1^ with a step size of 2 cm^−1^ on cells incubated with PA-d31 in the presence or absence of the SCD inhibitor CAY10566. The MIP images of the cells from SCD-active and SCD-inhibited groups are shown in **Fig. 2A**. Asymmetrically reweighted penalized least squares (arPLS) analysis (*27*), a baseline correction algorithm, was applied to remove the intracellular water background, allowing for accurate evaluation of the real intensity values of the targeted peak. The arPLS validation is described in **Supplementary Note 1**. Briefly, arPLS baseline correction was applied to remove the intracellular water background from the hyperspectral MIP spectra. The corrected PA-d31 substrate spectrum showed a flat baseline in the region of interest from 2230 cm^−1^ to 2260 cm^−1^. Moreover, the intracellular spectra from SCD-inhibited group also matched up with that of substrate in water, suggesting arPLS enables accurate detection of the 2246 cm^−1^ peak without interference (**Fig. S3**). The following intracellular spectra presented in this work were processed using arPLS baseline correction unless otherwise stated. The MIP spectra (**Fig. 2B**) obtained from the hyperspectral images of the inhibitor-free and inhibitor-treated groups showed clear signaling differences at 2246 cm^−1^, reflecting altered SCD activity. Statistical analysis (**Fig. 2C)** of the normalized 2246 cm^−1^ intensity by the 2200 cm^−^1 peak intensity showed a significant difference between the SCD-active and SCD-inhibited groups, supporting that the 2246 cm^−1^ peak is linked to SCD activity and strongly reduced upon SCD inhibition. To ensure an objective analysis, an independent region of interest (ROI) selection workflow was used (**Fig. S4**). Briefly, a mask was generated from the C-D channel corresponding to newly synthesized LDs to identify ROIs, which were then applied to the entire hyperspectral dataset to extract spectra for statistical analysis.

**Figure 2.**
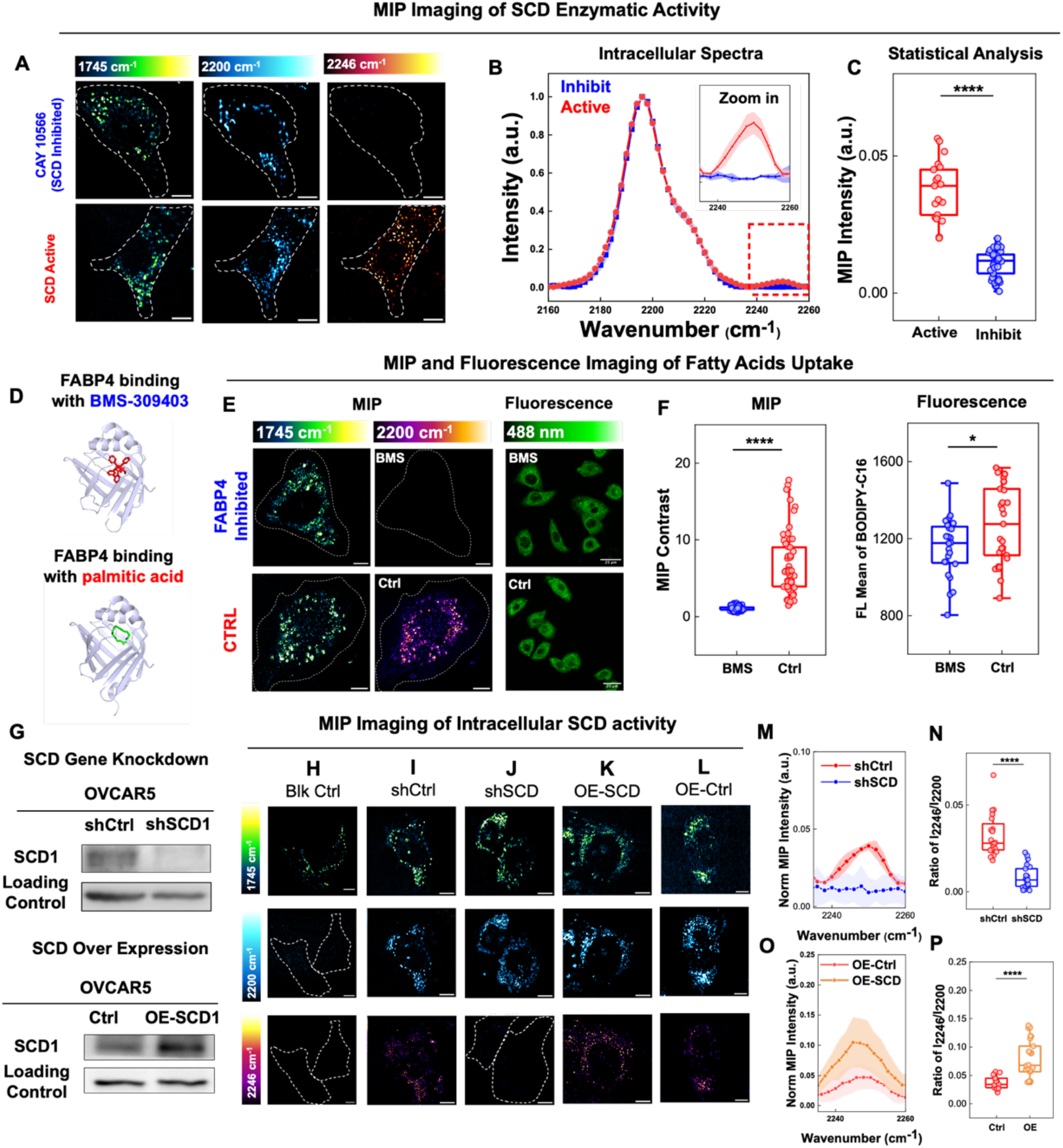
Measuring intracellular SCD activity. **(A)** MIP images of SCD-inhibited (top) and SCD active cells (bottom) at 1745 cm^−1^, 2200 cm^−1^, and 2246 cm^−1^. Scale bar, 10 μm. **(B)** Hyperspectral MIP spectra of the SCD-inhibited and the SCD-active group. **(C)** I_2246_/I_2200_ ratio in the SCD-inhibited and the SCD-active group. Active: n = 21. Inhibit: n = 41, **** p = 1.8×10^−20^. **(D)** Reported structures of FABP4 bound to palmitic acid (PDB code: 2HNX) (*38*) and BMS-309403 (PDB code: 2NNQ) (*39*). **(E)** MIP images of FABP4-inhibited cells (top) and control cells (bottom) at 1745 cm^−1^ and 2200 cm^−^1. Scale bar, 10 μm. Confocal fluorescence images of PA BODIPY in FABP4 inhibited (top right) and control cells (bottom right). Scale bar, 25 μm. **(F)** Statistical analysis of normalized MIP intensity (left) and PA-BODIPY fluorescence intensity (right) in FABP4 inhibited and control cells. BMS concentration: 2.5 μM. For MIP, BMS: n = 96. Ctrl: n = 60, **** p = 4.5×10^−44^. For PA-BODIPY, n = 16 cells, each point representing one cell, * p = 0.022. **(G)** Western blot of SCD1 in shCtrl, shSCD, OE-SCD, and control cells. **(H-L)** MIP images of blank control **(H)**, shCtrl **(I)**, shSCD **(J)**, SCD overexpression **(K)**, and control cells **(L)** at 1745 cm^−1^, 2200 cm^−1^, and 2246 cm^−1^. Scale bar, 10 μm. **(M)** Normalized MIP spectra of shCtrl and shSCD cells. **(N)** I_2246_/I_2200_ ratio in shCtrl and shSCD cells. shCtrl: n = 22, shSCD: n = 23, **** p = 1.2×10^−9^. **(O)** Normalized MIP spectra from OE-SCD and control cells. **(P)** I_2246_/I_2200_ ratio in OE-SCD and control cells. OE: n = 28. Ctrl: n = 22, **** p = 1.1×10^−5^.

To further validate that the desaturated signal is from LDs, we conducted fluorescence-detected MIP (F-MIP) imaging (*31*). In F-MIP, IR absorption modulates the fluorescence intensity of thermosensitive dyes in labeled organelles (*32*), the localization of MIP signal can be suggested. OVCAR5 cells were stained with LipiRed and treated with PA-d31 under the same experimental condition. As shown in **Fig. S5**, fluorescence imaging precisely revealed the locations of intracellular LDs. The corresponding F-MIP spectra collected from LDs reflect a clear peak at 2246 cm^−1^, consistent with the scanning MIP results. To verify that the newly generated peak specifically arises from SCD activity rather than other desaturases, we used SC-26196 to inhibit fatty acid desaturase 2 (FADS2), a major desaturase involved in cellular FA desaturation. As shown in **Fig. S6**, the 2246 cm^−1^ peak remains in the presence of SC-26196, verifying that the newly generated peak at 2246 cm^−1^ is from SCD activity rather than FADS2 activity.

Since PA-d31 serves as a PA analogue to mimic the metabolic process of PA, we examined whether it is taken up through native FA metabolic pathways. OVCAR5 FA uptake is facilitated by transporter proteins such as fatty acid binding proteins (FABPs) on the cell membrane (*8, 33–35*). Thus, we compared the PA-d31 uptake with or without FABP inhibitor treatment. BMS-309403 was used as the selected FABP inhibitor as it can occupy the FA-binding pocket of FABP, demonstrating significant inhibitory effects and blocking FA uptake in ovarian cancer cells (*1, 36, 37*). **Fig. 2D** shows the binding structures of FABP4 with the BMS-309403 inhibitor (top) and palmitic acid (bottom). As shown in **Fig. 2E**, MIP images of the cells in inhibitor-free group (Ctrl) exhibited strong signal at 2200 cm^−1^, corresponding to the CD_2_ asymmetric stretch, which serves as the reference of PA-d31 uptake. However, the FABP-inhibited group using BMS did not show detectable C-D intensity at 2200 cm^−1^, indicating effective inhibition of PA-d31 uptake, confirmed by the corresponding MIP spectra **(Fig. S7)**. Further statistical analysis **(Fig. 2F**, left) also confirmed the significant differences between the BMS and control group. These results indicate that PA-d31 uptake involves FABP-mediated FAs transport in ovarian cancer cells, and that inhibition of FABPs effectively suppresses PA-d31 uptake, supporting the use of PA-d31 as a physiologically relevant probe for monitoring SCD activity. We further compared the uptake of PA-BODIPY, a conventional fluorophore-labeled FA probe, between control and FABP-inhibited groups under the same conditions used in the MIP experiments. Bright intracellular BODIPY signals were observed in both groups **(Fig. 2E)**. In contrast to the pronounced change in MIP intensity, fluorescence intensity displayed only a modest yet statistically significant difference **(Fig. 2F**, right) between the FABP-inhibited and control groups. Similar results **(Fig. S8)** were obtained using BODIPY-C12. These results highlight the advantage of deuterated probe for faithfully reporting FABP-dependent FA uptake.

To validate the reliability of MIP imaging for monitoring intracellular SCD activity, we further explored two control conditions: (**i**) genetic knockdown (shSCD), and (**ii**) genetic overexpression (OE-SCD). The shSCD and OE-SCD generations were validated by Western blotting (WB) **(Fig. 2G)**. As shown in **Fig. 2H**, MIP imaging of the cells without PA-d31 treatment is free of C-D signal at both 2246 and 2200 cm^−1^, confirming minimum background interference. In **Fig. 2I**, cells in knockdown control (shCtrl) group exhibit obvious MIP signal at 2246 cm^−1^ and 2200 cm^−1^, suggesting that PA-d31 underwent desaturation during the PA-d31 incubation. However, cells in the shSCD group (**Fig. 2J**) did not show detectable signal from D-C=C-D at 2246 cm^−1^, indicating the SCD activity is strongly suppressed due to the gene knockdown. In **Fig. 2K**, OE-SCD cells served as the positive control and showed a stronger signal at 2246 cm^−1^ compared with cells with normal SCD expression (**Fig. 2L**), indicating increased SCD activity upon SCD overexpression.

We then compared hyperspectral MIP profiles across these conditions (**Fig. 2M** and **O**). The C-D MIP spectral profiles were normalized to the 2200 cm^−1^ peak. The shCtrl group (**Fig. 2M** and **N**, red) and SCD regular expression groups (**Fig. 2B and C**, red; **Fig. 2O** and **P**, red) exhibited comparable peak intensities at 2246 cm^−1^, indicating consistent and detectable FA desaturation under endogenous SCD expression. In contrast, the characteristic 2246 cm^−1^ peak was strongly reduced in the shSCD group (**Fig. 2M** and **N**, blue), consistent with the result in the pharmacological inhibition group (**Fig. 2B** and **C**, blue). The OE-SCD group cells displayed a significantly higher 2246 cm^−1^ peak intensity (**Fig. 2O** and **P**, orange) than the cells expressing endogenous levels of SCD (**Fig. 2O and P**, red), further supporting the assignment of the 2246 cm^−1^ peak to the SCD activity.

### Evaluation of Enzymatic activity by the Apparent SCD Activity Index

To enable accurate comparison of apparent intracellular SCD activity across different biological conditions, we established a readout that accounts for the variations in FA uptake changes during product formation, as shown below. Here, we define ***K***_***app***_as the apparent SCD activity index. ***V***_***max***_ represents the maximum reaction rate. ***K***_***m***_(***SCD***) is the Michaelis constant of SCD. [***P***] and **[*S***] represent the concentrations of the product and substrate of the SCD enzymatic reaction, respectively, while ***t*** represents the incubation time. Based on Michaelis-Menten equation and our experimental results, we derived **Eq. 1**, which supports the use of I_2246_/I_2200_ ratio as a readout of the apparent SCD activity index. The detailed mathematical derivation and theoretical discussion are provided in **Supplementary Note 2 (Eqs. S-1** to **S-8)**.

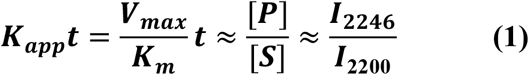

To ensure accurate peak intensity measurements from the complex intracellular environment, arPLS baseline correction was applied to remove the broad intracellular water background (*27*). As validated in **Supplementary Note 1**, this arPLS workflow reliably removed the water baseline from control substrate spectra without introducing artificial features in the spectral region of interest.

We experimentally tested the robustness of this readout by comparing cells treated with either 25 μM or 50 μM at the same incubation time (**Fig. S9A**). Although the cells incubated with 25 μM PA-d31 exhibited lower total FA uptake than those in the 50 µM group, the calculated I_2246_/I_2200_ ratio showed no statistically significant difference between the two substrate concentrations (**Fig. S9B**). The results confirmed our theory in **Supplementary Note 2**, within the tested concentration range, the I_2246/_I_2200_ ratio is not significantly affected by the intracellular substrate concentration and remained consistent at the same incubation time.

We also examined the time-dependent change of the intracellular I_2246_/I_2200_ ratio (**Fig. S10A** to **E**). The 2246 cm^−1^ peak becomes detectable after 6 h, I_2246_/I_2200_ ratio showed an approximately linear increase between ~8 h and ~24 h and approached a near-plateau after ~31 h of incubation (**Fig. S10F**).

### Elevated SCD activity in cisplatin treated cells

Next, we examined the SCD activity alteration in the cisplatin treated OVCAR5 cells (**Fig. 3A**) based on the established apparent SCD activity index. Cisplatin, a first-line chemotherapeutic agent for ovarian cancer, induces DNA damage and triggers extensive metabolic reprogramming (*40*). Previous studies have reported dysregulation of FA metabolism following cisplatin treatment (*41–43*), suggesting that SCD activity may be reprogrammed as part of the metabolic response to cisplatin.

**Figure 3.**
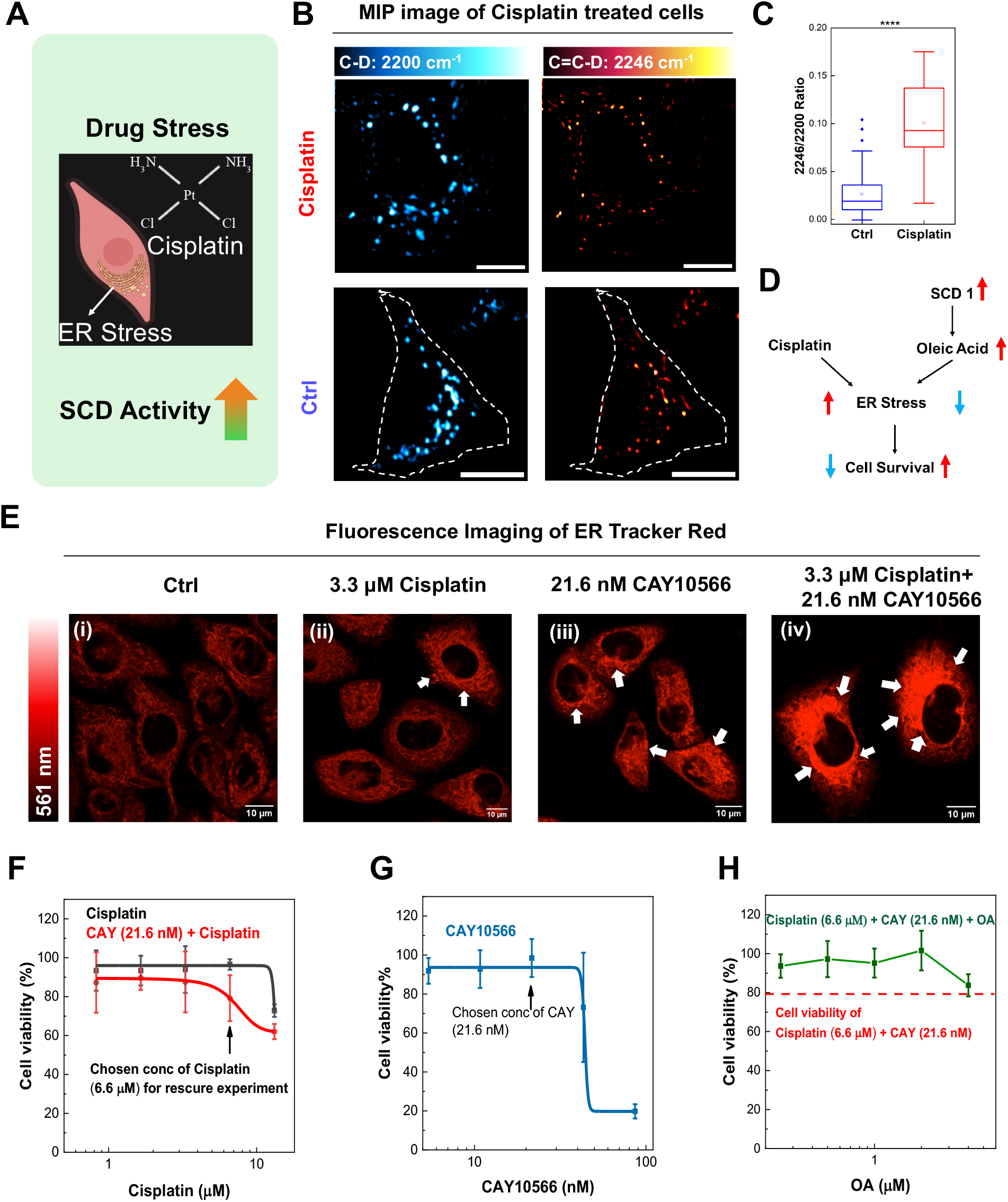
Cisplatin-induced endoplasmic reticulum (ER) stress increases SCD activity in OVCAR5 cells. **(A)** Schematic summary of SCD activity increased in cisplatin treated conditions. Created in BioRender. He, H. (2026) https://BioRender.com/t1ssolv **(B)** Single color images of cisplatin treated and control groups at 2200 cm^−1^ and 2246 cm^−1^. Cisplatin concentration: 3.3 μM, PA-d31 concentration: 50μM. Scale bar, 10 μm. **(C)** Statistical analysis of SCD activity index in control (blue) and cisplatin treated (red) group. Cisplatin: n = 35, Ctrl: n =44. **** p = 1.5×10^−8^. **(D)** Proposed mechanism of SCD activity promoting cell viability under cisplatin treatment. **(E)** Confocal fluorescence images of OVCAR5 cells labeled with ER tracker red. Scale bar, 10 μm. White arrows indicate regions of ER stress. Cisplatin concentration: 3.3 μM, CAY 10566 concentration: 21.6 nM. **(F-H)** Cell viability of OVCAR5 cells under different treatment conditions: cisplatin with or without CAY **(F)**, CAY alone **(G)**, and combined cisplatin, CAY, and OA treatment **(H)**. Cisplatin concentration: 6.6 μM, CAY 10566 concentration: 21.6 nM. OA concentration: 50 μM

As shown in **Fig. 3B**, clear MIP signal at 2246 cm^−1^ was observed in both cisplatin treated and control groups, confirming the conversion of PA-d31 to its corresponding unsaturated product. As shown in **Fig. 3C**, the SCD activity index within LDs showed a statistically significant increase following cisplatin treatment, suggesting that SCD activity is increased to cope with cisplatin induced stress.

Cisplatin treatment has been reported to induce endoplasmic reticulum (ER) stress (*44, 45*). We hypothesize that SCD activity may alleviate ER stress and thereby promote cell survival by generating MUFA including OA and palmitoleic acid (POA) (**Fig. 3D**). To test this hypothesis, we used confocal fluorescence imaging with an ER tracker to visualize ER morphology under four conditions: **(i)** control, (D) cisplatin treatment, **(iii)** SCD inhibition alone, and **(iv)** SCD inhibition combined with cisplatin treatment (**Fig. 3E)**. The control group exhibited a smoothed and typical interconnected ER network. Cisplatin treatment alone showed mild ER stress. SCD inhibition alone also caused mild ER alterations. However, the combination of SCD inhibition and cisplatin treatment resulted in disrupted ER morphology. These results suggest that SCD activity plays a protective role in maintaining ER homeostasis under cisplatin induced stress, likely through the production of MUFA.

Consistent with the ER morphology observations, MTT cell viability assays further supported the protective role of SCD activity under cisplatin-induced stress. As shown in **Fig. 3F**, inhibition of SCD by CAY10566 significantly reduced cell viability under cisplatin treatment compared with cisplatin alone, suggesting that SCD activity supports cell survival under cisplatin induced stress. To exclude intrinsic toxicity of the inhibitor, we examined the effect of CAY10566 alone. At the selected concentration (21.6 nM), CAY10566 did not significantly affect cell viability (**Fig. 3G**). Finally, supplementation with MUFA restored cell viability in cells treated with both cisplatin and the SCD inhibitor (**Fig. 3H**), supporting that SCD activity products contribute to cell survival under cisplatin induced stress.

Together, these results suggest that SCD activity is increased in cisplatin treated ovarian cancer cells. SCD may promote cancer cell survival during cisplatin treatment by generating MUFA and reducing ER stress.

### Elevated SCD Activity and Membrane Incorporation of SCD Products in Stem Cells

Aldehyde dehydrogenase-positive (ALDH+) cells in ovarian cancer are commonly used as an ovarian cancer stem cell (CSCs) enriched population and are associated with tumor growth, therapy resistance, and distinct metabolic features (*46, 47*). We asked whether SCD activity is reprogrammed in CSCs **(Fig. 4A and G)**. Using MIP hyperspectral imaging, we measured the SCD activity in ALDH+ (**Fig. 4B and C)** and ALDH-(**Fig. 4H and I**) groups. Both groups showed clear signal in the 2246 cm^−1^ channel (**Fig. 4C** and **I**). We also directly visualized the subcellular distribution of SCD products in CSCs with hyperspectral MIP imaging.

**Figure 4.**
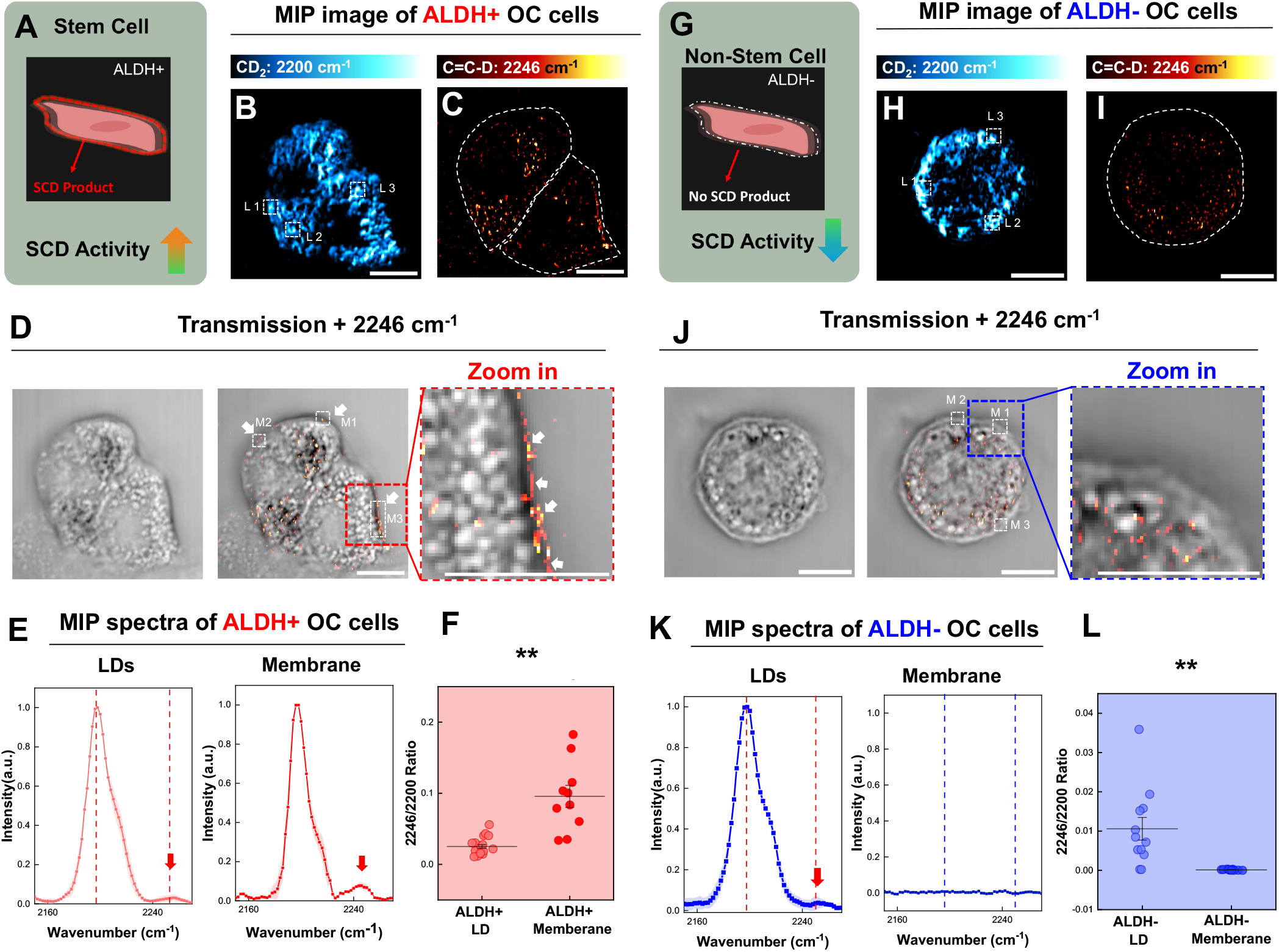
SCD activity increased in ALDH+ group comparing with ALDH− group. **(A, G)** Schematic summary of SCD activity increased in CSCs and SCD product participates in CSCs membrane biosynthesis. Created in BioRender. He, H. (2026) https://BioRender.com/t1ssolv **(B-C)** MIP imaging of ALDH+ OC cells at 2200 cm^−1^ and 2246 cm^−1^. PA-d31 concentration: 50 μM. Scale bar, 10 μm. **(D)** Overlap of MIP imaging of ALDH+ OC cells at 2246 cm^−1^ channel and the corresponding transmission channel. Scale bar, 10 μm. **(E)** MIP hyperspectral spectra from ALDH+ OC cells in the selected area in **(B)** and **(D)**. **(F)** SCD activity index comparison of lipid droplets and membrane area within ALDH+ group. LD: n = 21. Membrane: n = 19. ** p = 0.0013. **(H) (I)** MIP imaging of ALDH− OC cells at 2200 cm^−1^ and 2246 cm^−1^. Scale bar, 10 μm. **(J)** Overlap of MIP imaging of ALDH− OC cells at 2246 cm^−1^ channel and the corresponding transmission channel. Scale bar, 10 μm. **(K)** MIP hyperspectral spectra from ALDH− OC cells in the selected area in **(H)** and **(J)**. **(L)** SCD activity index comparison of lipid droplets and membrane area within ALDH− group. LD: n =14. Membrane: n = 14, ** p = 0.0039.

Spatially, both ALDH+ and ALDH− groups exhibited strong C-D signals in LDs regions (**Fig. 4B and H**), reflecting PA-d31 uptake and subsequent incorporation into cellular lipid pools. However, only ALDH+ cells displayed a significant signal in 2246 cm^−1^ channel at the cell boundary, likely the plasma membrane (white arrows in **Fig. 4D**), whereas this membrane-associated signal was absent in ALDH− cells (**Fig. 4J**), suggesting that some SCD products is used for membrane lipid remodeling instead of being stored in LDs in the ALDH+ group. Hyperspectral analysis with arPLS baseline correction further supported this observation. A distinct C-D stretching peak was detected in the membrane regions of ALDH+ cells but absent in ALDH− cells (**Fig. 4E** and **K**). This spatial distribution highlights the functional significance of SCD activity in CSCs, not only in lipid storage but also in remodeling membrane composition, likely contributing to membrane fluidity (*48*).

Quantitatively, within the ALDH+ group, SCD activity index in cell boundary regions is higher than that in LD regions (**Fig. 4F**), suggesting the membrane region has a higher level of newly synthesized SCD product. In contrast, the ALDH− group showed SCD product signals predominantly in LD regions (**Fig. 4L**). Statistical analysis across the two groups (**Fig. S11**) further showed that ALDH+ cells had a higher overall SCD activity index than ALDH− cells, likely reflecting the greater aggressiveness of the ALDH+ population. Together, these results show elevated SCD activity in CSCs and suggest that SCD products may contribute to membrane biosynthesis.

### Reduced SCD Activity under Oleic Acid (OA) Supplementation

To examine whether OVCAR5 cells adjust SCD activity under exogenous OA supplementation (**Fig. 5A**), we treated the cells with either PA-d31 alone or with OA combined with PA-d31 (**Fig. 5B**). Imaging results revealed the presence of larger LDs in the experimental group (**Fig. 5B** top) compared to the ones in control group (**Fig. 5B** bottom), likely resulting from the higher total FA concentration. Based on the established apparent SCD activity index, statistical analysis (**Fig. 5C**) showed a significant decrease in SCD activity index in the MUFA-supplemented group, indicating reduced SCD activity in response to intracellular MUFA enrichment. These results suggest that ovarian cancer cells dynamically adjust SCD activity according to intracellular MUFA flux, thereby maintaining SFA-USFA balance (*7*). We next examined whether SCD1 expression is downregulated when the SCD activity decreases. Immunofluorescence (IF) experiments were performed under the same conditions as the MIP imaging experiments to assess SCD1 protein levels, with nuclear stain by DAPI **(Fig. 5D)**. As shown in **Fig. 5E**, the IF signal exhibited comparable intensity (not significant) between the control and MUFA-supplemented group, indicating that SCD1 expression levels remained unchanged while SCD activity decreased. These results demonstrate that bond-resolved MIP imaging enables direct functional measurement of intracellular SCD activity alteration, which cannot be inferred from protein expression levels alone.

**Figure 5.**
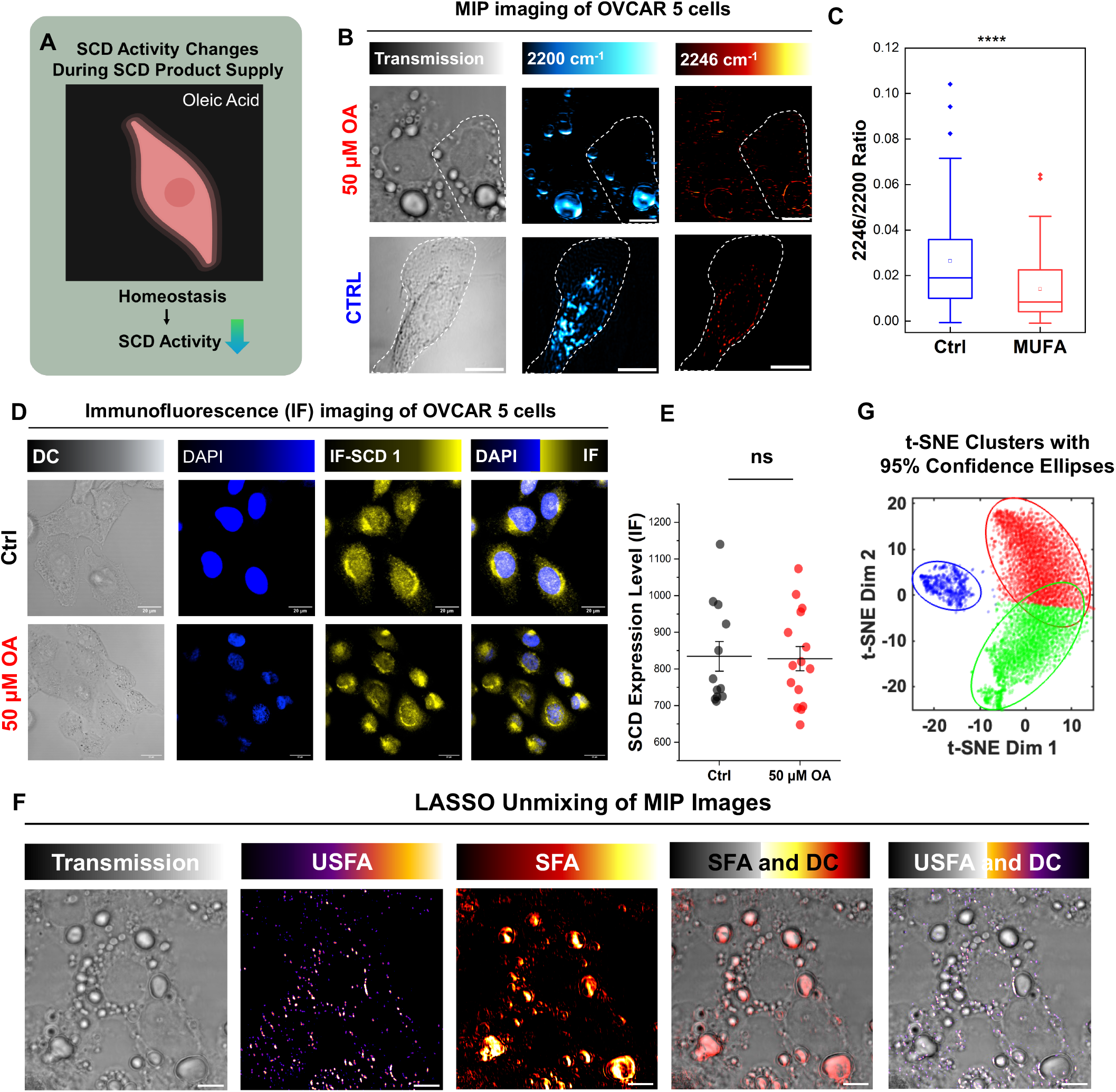
SCD activity decreases under conditions of OA supply and SCD product distribution. **(A)** Schematic summary of decreased SCD activity during OA supplementation. Created in BioRender. He, H. (2026) https://BioRender.com/t1ssolv **(B)** Single color images of PA-d31 only (Ctrl), PA-d31 and OA co-treated cells (50 μM OA) at D-C=C-D and C-D and corresponding transmission images. OA treated concentration: 50 μM, PA-d31 concentration: 50 μM. Scale bar, 10 μm. **(C)** Statistical analysis of OA-treated and control group. Ctrl: n = 46. OA: n = 90 *** p = 9.6×10^−5^. **(D)** Immunofluorescence images of SCD1 expression in PA-d31 and OA treated cells with DAPI staining. OA treated concentration: 50 μM.Scale bar, 20 μm. **(E)** SCD1 expression level comparison between PA-d31 and OA treated groups in **(D)**, n = 15. ns, p = 0.072. **(F)** LASSO unmixing of MIP imaging of OA and PA-d31 supplied cells. Scale bar, 10 μm. **(G)** t-SNE analysis of MIP hyperspectral imaging with 95% confidence ellipses. Saturated fatty acid (SFA, red), water background (BG, green), unsaturated fatty acid (USFA, blue).

### Spatial Distribution of SCD Products

Because the deuterium-labeled FA is incorporated into LDs in both saturated and unsaturated forms, we questioned whether the desaturation product is evenly distributed to LDs. To differentiate the SFA and USFA spatial distribution, we introduced the least absolute shrinkage and selection operator (LASSO) (*49*)to unmix the three major components in the cell silent window (**Fig. S12**): **(i)** SFA, **(ii)** USFA, **(iii)** water background (BG). Notably, the single-color images at 2200 cm^−1^ and 2246 cm^−1^ present the signal of total C-D and D-C=C-D, respectively. They cannot spatially reveal both SFA and USFA species. LASSO unmixing was therefore employed to resolve these components based on their distinct full spectral profiles. To validate the ability to unmix SFA and USFA, we applied LASSO unmixing to both SCD-active and SCD-inhibited groups (**Fig. S13**). USFA components were detected in cells from the SCD-active group but were absent in the SCD-inhibited group, indicating that the LASSO unmixing method performed reliably in our system.

As shown in **Fig. 5F**, the spatial distribution of SFA and USFA was successfully resolved using LASSO. To improve spatial visualization, the extracted signals were overlaid with the transmission channel. The results clearly demonstrate that desaturation products predominantly localize to the LD surface, whereas SFAs are concentrated in the LD core. This result is repeatable in **Fig. S14**. We further examined the 2246 cm^−1^ peak profile of an enlarged LD in the dashed region of **Fig. S15A** and **B**. A clear 2246 cm^−1^ signal was observed on the LD surface in **Fig. S15C** and **D**, with consistent validation from the corresponding MIP spectra (**Fig. S15E and F**). This analysis revealed that the SCD product was not uniformly distributed throughout the LD, suggesting a core-shell spatial organization with higher SCD product content near the LD shell.

To independently check the feature separation observed in the LASSO analysis, we further applied t-distributed stochastic neighbor embedding (t-SNE), an unsupervised nonlinear dimensionality reduction method for data visualization (*50*). Our result (**Fig. 5G**) showed that the three clusters are clearly separated with minimal overlap, demonstrating that the three components identified by LASSO have spectral signatures that can also be distinguished by an unsupervised method. These results reveal a spatial organization of SFA and USFA within LDs. Because the spatial resolution of the current MIP measurement is approximately 230 nm, intra-droplet spatial differences are difficult to reliably resolve in small lipid droplets, especially those in control cells with diameters often below or close to this resolution limit. The treatment condition with OA and PA co-treated generated larger LDs, allowing the spatial distribution of the product associated signal to be compared within individual droplets.

## Discussion

From a therapeutic perspective, our findings highlight SCD as a potential target in ovarian cancer cells. While inhibition of SCD alone exerted minimal effects on cell viability, SCD inhibition markedly sensitized cells to cisplatin treatment, an effect that could be reversed by exogenous MUFA supplementation. This indicates that SCD activity becomes critical under chemotherapeutic stress, where membrane homeostasis and lipid adaptability are required for survival.

Technically, we note that Raman spectroscopy offers an excellent approach for mapping intracellular metabolites. In particular, stimulated Raman scattering (SRS) microscopy is an exceptionally powerful and high-speed vibrational imaging modality for tracking cellular metabolism with small vibrational tags. Hyperspectral SRS has been used to map deuterium labeled fatty acid uptake in cells (*51*), glucose metabolism in cells (*52*) and animals (*53*). Recently, high content hyperspectral SRS imaging also combined with LASSO analysis was demonstrated for mapping multiple metabolites (*1*). However, direct SRS imaging of SCD-catalyzed fatty acid desaturation through the newly generated D-C=C-D has not been reported.

Previous studies via SRS microscopy have enabled imaging of lipid metabolism, including overall lipid unsaturation (*9*). However, overall lipid unsaturation mainly serves as a metabolic biomarker and these studies cannot distinguish newly synthesized unsaturated components from those acquired through exogenous uptake, neither giving any information about enzymatic reaction.

To explore the complementary capabilities of SRS and MIP for resolving the SCD product associated peak, we performed comparative hyperspectral measurements by both modalities under the same treatment condition using PA-d31 treated and OA-d34 treated cells (**Fig. S16**). Both samples are expected to show a product peak at 2246 cm^−1^. In the OA-d34 treated cells, this feature was detectable by both MIP and SRS. However, in the PA-d31 treated cells, although the newly generated 2246 cm^−1^ product associated peak was detected by MIP, it remained challenging to distinguish this weak feature from the strong adjacent C-D background envelope under our current SRS experimental settings. This difference is context dependent and likely reflects the complementary contrast mechanisms of the two modalities rather than a generalized limitation of either method. SRS provides outstanding sensitivity for detecting the bulk C-D labeled lipid pool based on Raman polarizability. In this specific case, the amount of desaturation product generated by the enzymatic reaction is limited, making the 2246 cm^−1^ peak intrinsically weak and embedded within the broad total C-D stretching. In contrast, MIP employs a photothermal readout to detect absorption, whereby the weak IR absorption of the minor product is converted into scattering modulation of the visible probe beam. This conversion is particularly effective within droplet-like structures, where scattering of the visible probe beam is magnified, thereby enhancing the overall detectability. These results support that SRS and MIP are complementary modalities: SRS excels in high-speed mapping of total lipid trafficking, whereas MIP offers a useful approach for resolving selected molecular transformations in complex intracellular environments.

In summary, using PA-d31 as a SCD activity reporter, we identified a distinct MIP signal at 2246 cm^−1^ corresponding to C=C bond formation due to SCD activity, a key metabolic feature of ovarian cancer reprogramming. Hyperspectral MIP imaging revealed elevated SCD activity in ovarian cancer cells under cisplatin-induced stress, suggesting a potential role of SCD products in reducing ER stress. In contrast, exogenous supplementation of MUFAs suppressed SCD activity without altering SCD expression levels, indicating feedback regulation of SCD activity. Furthermore, SCD activity was significantly increased in CSCs, where a portion of SCD products was incorporated into the plasma membrane. Spatial analysis further revealed heterogeneous lipid droplet organization, with SCD products enriched at the LD surface. Together, this work establishes bond-selective MIP imaging as a chemically specific strategy for visualizing intracellular enzymatic activity and lipid metabolic organization under diverse biological conditions.

## Materials and Methods

### Materials

BMS309403, cisplatin, and CAY10566 were purchased from Cayman Chemicals. SC26196 was purchased from Medchem Express.

### Cell culture

RPMI-1640 Culture medium (Thermofisher Scientific) with the addition of 10% (vol/vol) fetal bovine serum (FBS) and 1% (vol/vol) penicillin–streptomycin (P/S) was used for OVCAR5 (ATCC, HTB-161) cell culturing. Cells were cultured at 37°C in a humidified incubator with 5% CO_2_ supply.

### Cell line

OVCAR5 cells, SCD knockdown and SCD overexpression cell lines and their characterizations were kindly provided by Dr. Matei’s group. The cell lines were previously generated and characterized (*7*). The Western blot characterization presented in this study was performed as previously described (*7*).

### Cell treatment

PA-d31 or OA-d34 probe treatment group: the OVCAR5 cells were seeded with 10% FBS RPMI-1640 medium, and then 50 mM PA-d31 or OA-d34 stock solution was diluted with delipid RPMI-1640 medium for 24 h incubation, final concentration was 50 μM.

MUFA treatment group: the OVCAR5 cells were seeded with 10% FBS RPMI-1640 medium, and then 50 mM OA stock solution and 50 mM PA-d31 were diluted with delipid RPMI-1640 medium for 24 h incubation, final concentration was 50 μM.

Cisplatin treatment group: the OVCAR5 cells were seeded with 10% FBS RPMI-1640 medium, and then 3.3 mM cisplatin stock solution was diluted with 10% FBS medium for 24-hour incubation, final concentration is 3.3 μM. Following with PA-d31 treatment processes.

Inhibition group: all the inhibitors are pre-treated for 24 hours and co-treated with the deuterium probe to ensure the inhibition; concentrations are specified in the text and captions. Following with the cell treatment protocol as above.

Time-course: OVCAR5 cells were seeded on CaF_2_ slides overnight and then incubated with 50 μM PA-d31 in de-lipid RPMI-1640 medium for 3 to 48 h.

### Cancer stem cell isolation

Ovarian cancer stem cells were selected with high Aldehyde dehydrogenase (ALDH) activity. ALDH activity was measured by flow cytometry using an ALDEFLUOR assay kit (Stemcell Technologies, Cat#01700, Cambridge, MA, USA) following the manufacturer’s instructions and as described previously (*54, 55*). In brief, OVCAR5 cell pellets were suspended in the ALDEFLUOR assay buffer and incubated with the ALDEFUOR substrate, BODIPYaminoacetaldehyde (BAAA) for 45 minutes at 37 °C. The ALDH inhibitor, diethylamino benzaldehyde (DEAB) was used to block the baseline level of ALDH enzymatic activity. ALDH positive cells were determined based on a relative increase in FITC signals compared to DEAB treated cells, and sorted using the fluorescence activated cell sorting, BD FACSAria 5-Laser Sorter (BD Biosciences, San Jose, CA). A minimum of 10,000 events were analyzed using Cell Quest software (BD Biosciences, version 3.3, San Jose, CA, USA) for each experiment.

### MIP microscope

The system is built on an open microscope platform (SliceScope, Scientifica). A continuous-wave 532 nm laser (Samba, HUBNER Photonics) acted as the visible-light probe, while a tunable pulsed quantum cascade laser (MIRcat, Daylight Solutions) supplied the mid-infrared pump, covering wavelengths from 900 cm^−1^ to 2300 cm^−1^. A galvanometer mirror (Saturn 3B, ScannerMax) is used to perform laser scanning of the visible beam which was then focused by a water immersion objective (UPLANSAPO, 1.2 NA, 60×, Olympus). Meanwhile, the mid-infrared beam was synchronously scanned using another set of galvanometer mirrors (Saturn 5B, ScannerMax) and brought to the same focal point through a reflective objective (0.78 NA, 40×, PIKE Technologies). To reduce chromatic aberration during infrared beam scanning, an all-reflective relay composed of two concave mirrors with focal lengths of 200 mm and 500 mm was used. The microscope system was controlled using LabView 2020 software.

Intracellular measurements were performed using a counter-propagating beam geometry. The visible beam was delivered from above and focused onto the sample using a water-immersion objective (60×, 1.2 NA, UPlanSApo, Olympus), whereas the mid-infrared beam was delivered from below and focused using a reflective objective (40×, 0.78 NA, PIKE Technologies). For in vitro measurements, a co-propagating geometry was used, as shown in Fig. S2, in which both the visible and mid-infrared beams were delivered through the reflective objective.

### SRS microscope

The SRS system was lab-built, described in previously published work (*1*). The setup employed a femtosecond laser (InSight DeepSee, Spectra Physics, Santa Clara, CA, USA) operating at 80 MHz, which delivered two synchronized outputs: a tunable pump beam from 680 to 1300 nm and a fixed wavelength Stokes beam at 1040 nm. To probe the C-D stretching region, the pump beam was tuned to 847 nm. Modulation of the Stokes beam at 2.3 MHz was achieved using an acousto optic modulator (1205-C, Isomet). Following beam combination, both beams were temporally stretched using two SF57 glass rods, each 12.7 cm in length, before entering a laser scanning microscope. The optical powers of the pump and Stokes beams at the microscope input were adjusted to 30 mW and 200 mW, respectively. Light was focused onto the sample with a 60× water immersion objective (NA = 1.2, UPlanApo/IR, Olympus), and signal collection was performed with an oil condenser (NA = 1.4, U-AAC, Olympus).

### Fluorescence labeling and imaging

BODIPY-C12 (4,4-Difluoro-5,7-Dimethyl-4-Bora-3a,4a-Diaza-s-Indacene-3-Dodecanoic Acid) was purchased from ThermoFisher. Cells were incubated with BODIPY-C12 at a final concentration of 2 μM for 15 minutes at 37°C. Following incubation, cells were washed three times with PBS to remove excess dye. Imaging was performed after formalin fixation, using a confocal microscope (Olympus FV3000) with a 488 nm excitation laser source.

ER-Tracker Red (BODIPY TR Glibenclamide) was purchased from ThermoFisher. Cells were incubated with ER-Tracker Red at a final concentration of 1 μM for 30 minutes at 37 °C. Following incubation, cells were washed three times with PBS to remove excess dye. Imaging on a confocal microscope (Olympus FV3000) with a 561 nm excitation laser source.

For immunofluorescence imaging, samples were fixed with 4% paraformaldehyde, permeabilized with 0.1% Triton X-100, and blocked with 1% bovine serum albumin (BSA) in PBS. Cells were then incubated with anti-SCD1 primary antibody (ab236868, Abcam) followed by Alexa Fluor 488-conjugated secondary antibody. Nuclei were counterstained with 1 μM DAPI, imaging was performed on a confocal microscope (Olympus FV3000) with a 488 nm and 405 nm excitation laser source.

### FTIR spectroscopy

FTIR spectra were recorded using a Nicolet Nexus 670 spectrometer (Thermo Fisher Scientific) equipped with a diamond detector. The spectral resolution was set to 2 cm^−1^, and each spectrum was collected by performing 64 scans to ensure high signal-to-noise ratio and accuracy in the measurements.

### Sample preparation and calculation of limit of detection

The limit of detection (LOD) was defined as the minimum analyte concentration at which the MIP signal could be reliably distinguished from the background noise and was calculated as,

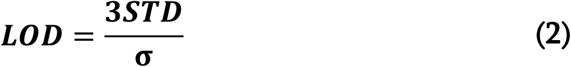

Here, ***STD*** is the standard deviation of the MIP intensity measured from the blank solvent at the region of interest, and **σ** represents the slope of the calibration curve relating analyte concentration to MIP signal intensity.

For LOD solution measurement, OA-d34 was first dissolved in DMSO and subsequently diluted with DMSO to final concentrations of 6.25, 12.5, 25, and 50 μM. Each solution was sandwiched between two CaF2 slides and measured using the co-propagating MIP configuration shown in **Fig. S1A**. The spectra were normalized to the solvent associated peak at 2035 cm^−1^. For thin-film measurements, PA-d31 and OA-d34 were separately dissolved in MeOH/DCM (v/v, 1:2) to prepare 50 mM stock solutions. The stock solutions were then mixed at appropriate ratios to obtain samples containing 0%, 10%, 50%, and 100% (molar ratio) OA-d34. Each mixture was drop-cast onto a CaF_2_ slide and allowed to form a thin film before MIP measurement, as shown in **Fig. S2A**.

### Statistical Analysis

Statistical analysis was performed using a two-tailed unpaired Welch’s t-test for comparisons between two groups. To keep the heterogeneity within one cell, multiple ROIs were selected from individual cells according to the subcellular structures of interest. Each data point shown in the scatter plots represents one ROI unless other statement. The numbers of analyzed ROIs (n) and independent cells (N) are indicated in the corresponding figure legends. The number of ROIs varied among cells depending on the number of identifiable subcellular structures. All eligible ROIs were included in the analysis. Data are presented as mean ± standard deviation. Statistical significance was defined as follows: ns, not significant; * p < 0.05; ** p < 0.01; *** p < 0.001; and **** p < 0.0001. Exact p values are shown in the captions. No samples, cells, or data points were intentionally excluded from the analyses.

### Baseline correction of MIP images

The data processing includes 4 steps: normalization and denoising, drift correction, background removal and mask encoding. The raw hyperspectral data is normalized and denoised by Block-Matching and 4D filtering (BM4D), a 3D version of BM3D (*56*). The BM4D algorithm applies grouping and collaborative filtering, stacking a 3D imaging block into a 4D data array for filtering, which attenuates the noise by utilizing the spatial and spectral correlation of the hyperspectral data. Next, Fast4Dreg (*57, 58*) was used to correct 3D drift in the data, by creating projections along multiple directions and estimating the drift using two-dimensional cross correlation. To estimate background interference, arPLS (*27*) was performed pixel by pixel to remove the water background in hyperspectral MIP data and extract the peak. The arPLS algorithm works by iteratively adjusting weights to fit a smooth baseline, while avoiding peaks in the spectrum. arPLS balances the signal above and below the baseline using a logistic function, refining the estimate until the correction is stable. Finally, to focus on lipids for statistical analysis, we generate intensity-based masks for segmentation. Morphological opening is applied to refine the selection, by removing small, isolated noise points and smoothing the boundaries of lipid regions, maintaining continuous and biologically relevant lipid structures. The spectra of lipid masks are then ready for further quantitative analysis and statistical assessments.

## Acknowledgments

The authors thank Haonan Lin for modifying the LASSO code.

## Funding

National Institutes of Health grant R33CA287046 (JXC) National Institutes of Health grant R35GM136223 (JXC) National Institutes of Health grant R01CA224275 (JXC DM) US Department of Veterans Affairs BX006012 (D.M)

## Data, Code, and Materials Availability

All data and code needed to evaluate and reproduce the results in the paper are present in the paper and/or the Supplementary Materials. This study did not generate new materials. Raw data and code from this study have been deposited in the research repository Zenodo under the following DOI: 10.5281/zenodo.21988655, accessible with the link: https://zenodo.org/records/21988655.

## Author contributions

Conceptualization: XT, JXC

Methodology: XT, HH, JXC

Investigation: XT, HH, JY, CVPD, YW, AI, ML, JA, GD, DD, QX

Visualization: XT, TX, HH

Supervision: JXC, DM, HH Writing—original draft: XT

Writing—review & editing: HH, JXC, DM

## Competing interests

All other authors declare they have no competing interests.

## Supplementary Materials

This PDF file includes:

Supplementary Note 1 and 2

Fig. S1 to S16

## Notes

### Competing Interest Statement

The authors have declared no competing interest.

### Summary of Updates

Added new results and theory framework.

