## Supplementary Materials for "Bio-orthogonal Mid-Infrared Photothermal Imaging Unveils Altered Stearoyl-CoA desaturase (SCD) Activity in Ovarian Cancer Cells"

Xinyan Teng *et al.*

**This PDF file includes:**

- Supplementary Note 1 and Note 2
- Figs. S1 to S16

- **Supplementary Note 1: arPLS Validation**

To validate that arPLS can effectively remove the background without introducing artifacts, we performed the control experiments. We validated the control substrate in water with the updated arPLS workflow. We measured pure PA-d31 in water (**Fig. S3A**), applied the arPLS baseline-correction procedure (**Fig. S3B**) to the data and used this arPLS-processed spectrum (**Fig. S3C**) rather than that of dry pure chemical as the spectral reference of substrate for the comparison with intracellular MIP spectra (**Fig. S3D**). Clearly, the substrate spectrum showed a flat baseline after arPLS correction, especially in the region of interest from 2230 cm^-1^ to 2260 cm^-1^. The arPLS processed intracellular MIP spectrum exactly matches up with that of PA-d31 in water (**Fig. S3D**). This result suggests that arPLS correction did not introduce artifacts into the spectrum in the spectral region used for 2246 cm^-1^ peak analysis.

- **Supplementary Note 2: Apparent SCD Activity Index Evaluation**

According to Michaelis-Menten equation, the SCD enzyme reaction rate can be described as:

|  | $\boldsymbol{v(SCD) =}\frac{\boldsymbol{V}_{\boldsymbol{max}}\boldsymbol{[S]}}{\boldsymbol{K}_{\boldsymbol{m}}\boldsymbol{+[S]}}$ | **(S-1)** |
| --- | --- | --- |

Here, $\boldsymbol{V}_{\boldsymbol{max}}$represents the maximum reaction rate, $\boldsymbol{K}_{\boldsymbol{m}}\boldsymbol{(SCD)}$ is the Michaelis constant. According to reported literature, $\boldsymbol{K}_{\boldsymbol{m}}\boldsymbol{(SCD)}$ is 10.5 μM (*59*). In our measurements, the limit of detection (LOD) for the C-D signal at 2200 cm^-1^ is 3.31 μM (**Fig. S1**). We didn’t observe detectable diffuse C-D substrate outside lipid droplets, suggesting that the concentration of the diffuse SCD substrate pool is much lower than reported $\boldsymbol{K}_{\boldsymbol{m}}\boldsymbol{(SCD)}$. Under the assumption of **[S]**$\boldsymbol{\ll}\boldsymbol{K}_{\boldsymbol{m}}\mathbf{(}\boldsymbol{SCD}\mathbf{)}$, the Michaelis-Menten equation can be simplified, and the reaction rate becomes approximately first-order with respect to substrate concentration:

|  | $\boldsymbol{v (SCD)}\boldsymbol{\approx}\frac{\boldsymbol{V}_{\boldsymbol{max}}}{\boldsymbol{K}_{\boldsymbol{m}}}\boldsymbol{[S]}$ | **(S-2)** |
| --- | --- | --- |

According to the time course measurement (**Fig. S10F**), the product formation remains approximately linear in the reaction time window from ~8 to ~24 h with $\boldsymbol{t}$ represents the incubation time,

|  | $\boldsymbol{[P]}\boldsymbol{\approx}\boldsymbol{vt}\boldsymbol{\approx}\frac{\boldsymbol{V}_{\boldsymbol{max}}}{\boldsymbol{K}_{\boldsymbol{m}}}\boldsymbol{[S]t}$ | **(S-3)** |
| --- | --- | --- |

[$\boldsymbol{P]}$ and **[**$\boldsymbol{S}\boldsymbol{]}$ represent the concentrations of the product and substrate of the SCD enzymatic reaction, respectively.

Therefore,

|  | $\frac{\boldsymbol{[P]}}{\boldsymbol{[S]}}\boldsymbol{\approx}\frac{\boldsymbol{V}_{\boldsymbol{max}}}{\boldsymbol{K}_{\boldsymbol{m}}}\boldsymbol{t}$ | **(S-4)** |
| --- | --- | --- |

Here, we defined the slope of linear window as $\boldsymbol{K}_{\boldsymbol{app}}$ which is also called the apparent SCD activity index. So,

|  | $\boldsymbol{K}_{\boldsymbol{app}}\boldsymbol{=}\frac{\boldsymbol{V}_{\boldsymbol{max}}}{\boldsymbol{K}_{\boldsymbol{m}}}$ | **(S-5)** |
| --- | --- | --- |

Because $I_{2246}$ is proportional to the abundance of the desaturation product, whereas $I_{2200}$ is proportional to the total abundance of deuterated species ($\left[ \boldsymbol{S} \right]\boldsymbol{+[P]}$). As $\boldsymbol{[P]}\boldsymbol{\ll}$ $\left[ \boldsymbol{S} \right]$. Thus,

|  | $\frac{\left[ \boldsymbol{P} \right]}{\left[ \boldsymbol{S} \right]\boldsymbol{+[P]}}\boldsymbol{\approx} \frac{\left[ \boldsymbol{P} \right]}{\left[ \boldsymbol{S} \right]}\boldsymbol{\approx}\frac{\boldsymbol{I}_{\boldsymbol{2246}}}{\boldsymbol{I}_{\boldsymbol{2200}}}$ | **(S-6)** |
| --- | --- | --- |

So,

|  | $\boldsymbol{K}_{\boldsymbol{app}}\boldsymbol{t=}\frac{\boldsymbol{V}_{\boldsymbol{max}}}{\boldsymbol{K}_{\boldsymbol{m}}}\boldsymbol{t}\boldsymbol{\approx}\frac{\boldsymbol{[P]}}{\boldsymbol{[S]}}\boldsymbol{\approx}\frac{\boldsymbol{I}_{\boldsymbol{2246}}}{\boldsymbol{I}_{\text{2200}}}$ | **(S-7)** |
| --- | --- | --- |

**So, for the same incubation time (e. g. t=24 h)**

|  | $\boldsymbol{K}_{\boldsymbol{app}}\boldsymbol{=}\frac{\boldsymbol{V}_{\boldsymbol{max}}}{\boldsymbol{K}_{\boldsymbol{m}}}\boldsymbol{\propto}\frac{\boldsymbol{[P]}}{\boldsymbol{[S]}}$ | **(S-8)** |
| --- | --- | --- |

Together, these results validate I_2246_/I_2200_ as a readout of apparent SCD activity index that accounts for the variations in fatty acid uptake changes during product formation. We therefore applied this readout to examine how SCD-catalyzed desaturation changes in response to different biological conditions.

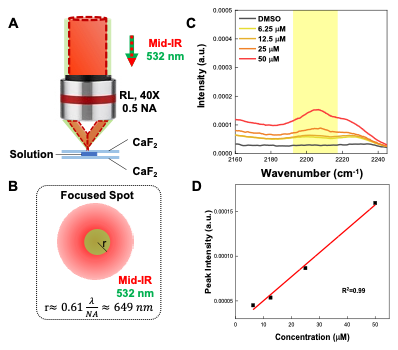

**Figure S1. Limit of detection measurement.**

(A) Schematic of co-propagation measurement method for solution samples.
(B) Mid-IR beam and visible beam focused spot on solution samples.
(C) MIP spectra of different concentration of OA-d34 solutions and DMSO.
(D) Linear fitting of peak of 2200 cm^-1^ in **(C)**.

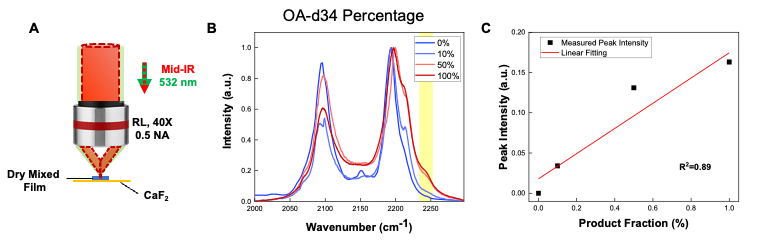

**Figure S2. MIP measurement of PA-d31/OA-d34 thin films using a co-propagating detection geometry.**

**(A)** Schematic of the co-propagating MIP detection geometry.

**(B)** MIP spectra of thin films containing mixed substrate and product standards with different OA-d34 fractions in PA-d31: 0%, 10%, 50%, and 100%.

**(C)** Linear fitting of the 2246 cm⁻¹ peak intensity as a function of the OA-d34 fraction.

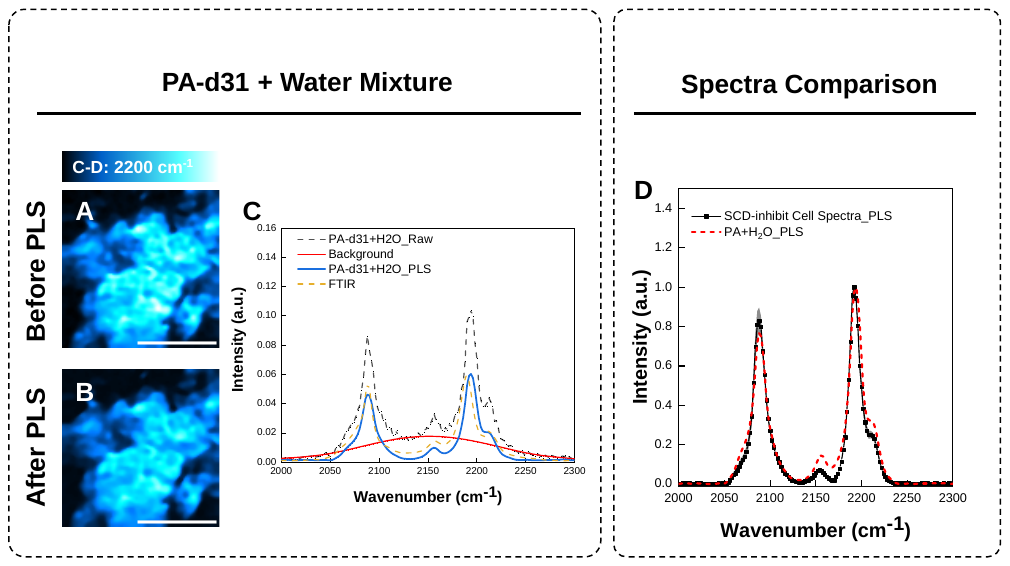

**Figure S3. arPLS processing and validation.**

**(A) (B)** MIP imaging of the mixture of PA-d31 pure chemical and water before and after arPLS. Scale bar: 20 μm.

**(C)** MIP spectra of pure PA-d31 immersed in water before (dashed black) and after (blue curve) arPLS baseline correction with FTIR reference of PA-d31 pure chemical (dashed orange).

**(D)** MIP spectra comparison of intracellular spectra in SCD-inhibited cell (black) and substrate spectra (red) after arPLS baseline correction.

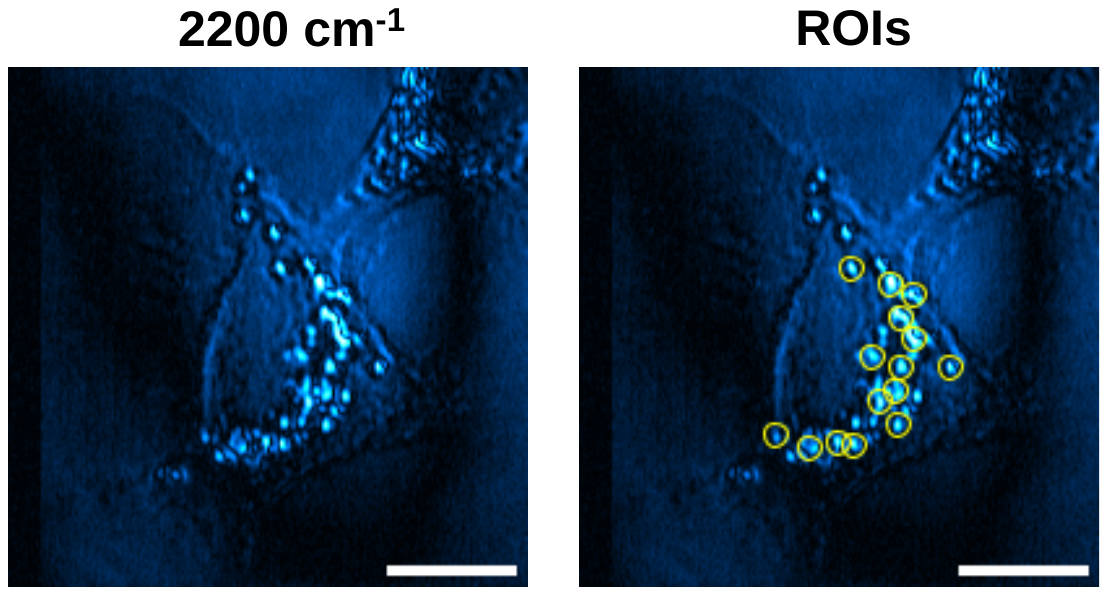

**Figure S4.** Workflow for ROIs selections in statistical analysis.

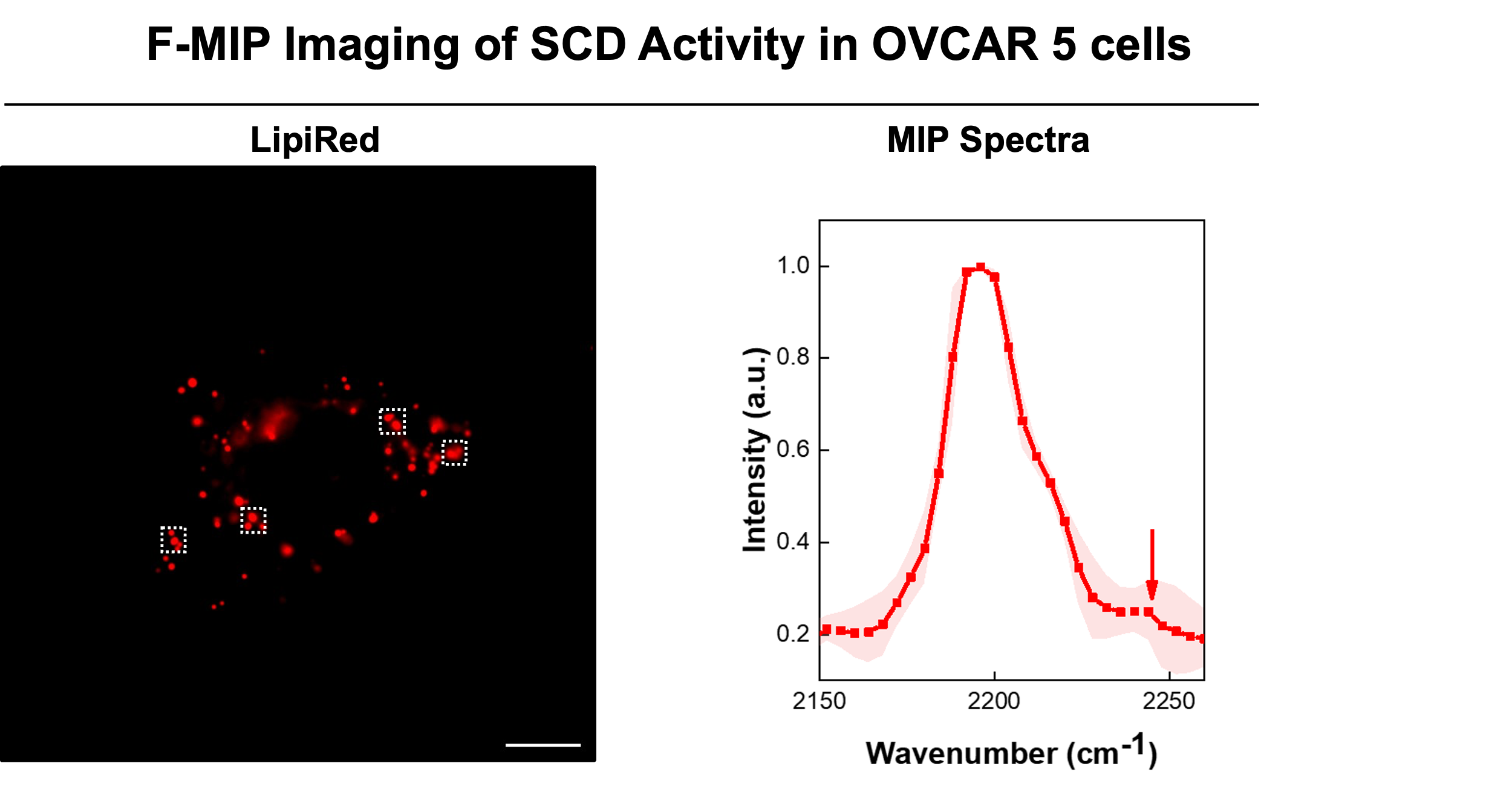

**Figure S5. F-MIP Imaging of PA-d31 treated OVCAR5 cells with lipid red staining.** Fluorescence imaging (left) and the corresponding MIP spectra (right) from the selected ROIs. Scale bar: 10 μm.

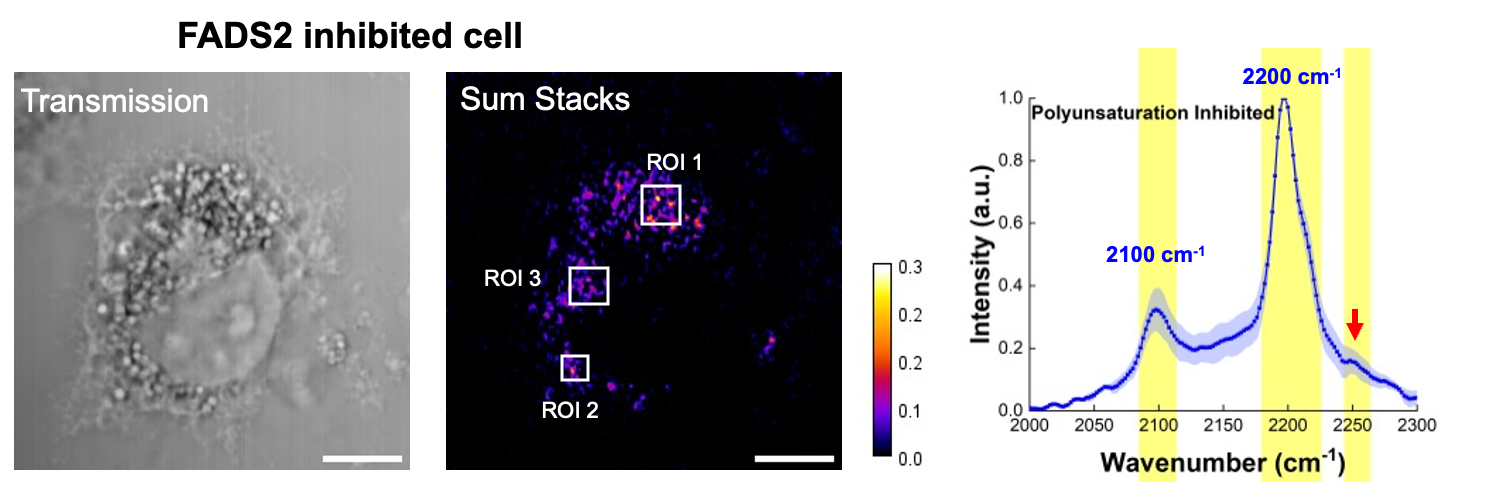

**Figure S6. MIP imaging of FADS2 inhibited cells.** MIP images and corresponding intracellular spectra of FADS2 inhibited cells. Scale bar: 10 μm.

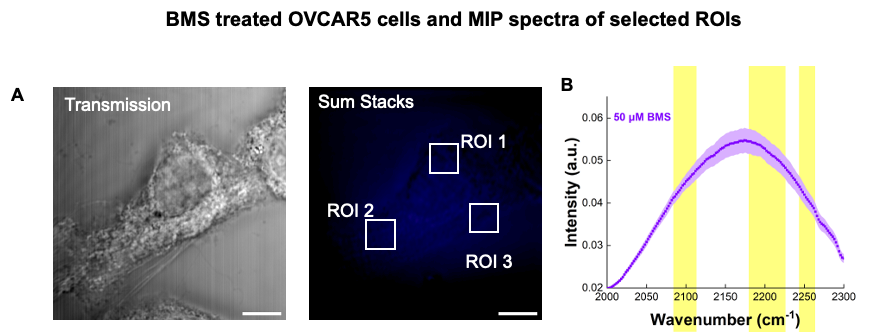

**Figure S7. FA uptake test of PA-d31 in OVCAR5 cells.** MIP images (A) and MIP spectra (B) of cells treated by BMS. Scale bar: 10 μm.

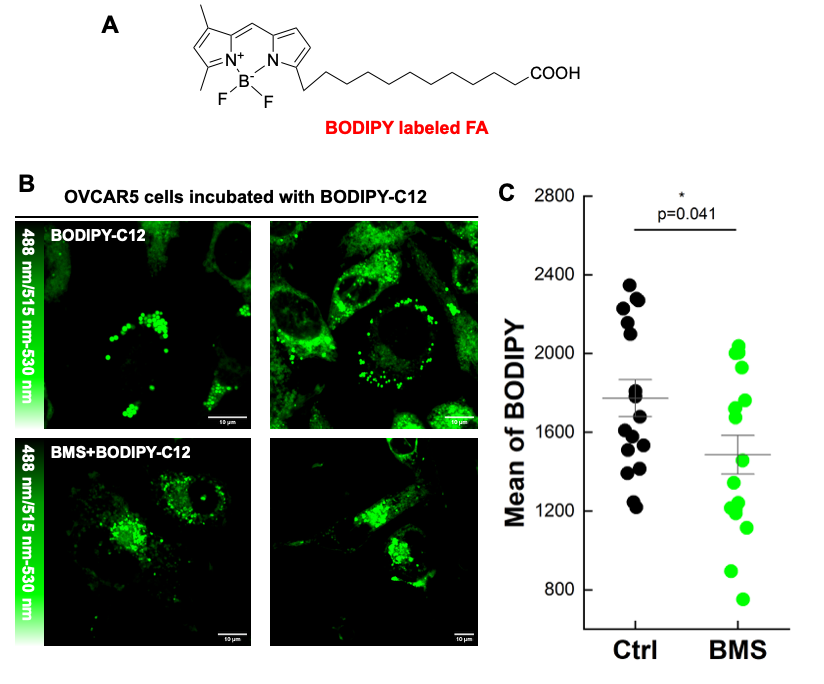

**Figure S8. FA uptake test of BODIPY-C12 in OVCAR5 cells.**

**(A)** BODIPY-labeled FA molecular structure.

**(B)** Fluorescence imaging of BMS inhibition and control groups. Scale bar, 10 μm.

**(C)** Statistical analysis of fluorescence intensity in **(B)**, * p = 0.041, N = 17 cells, with each data point represent one cell.

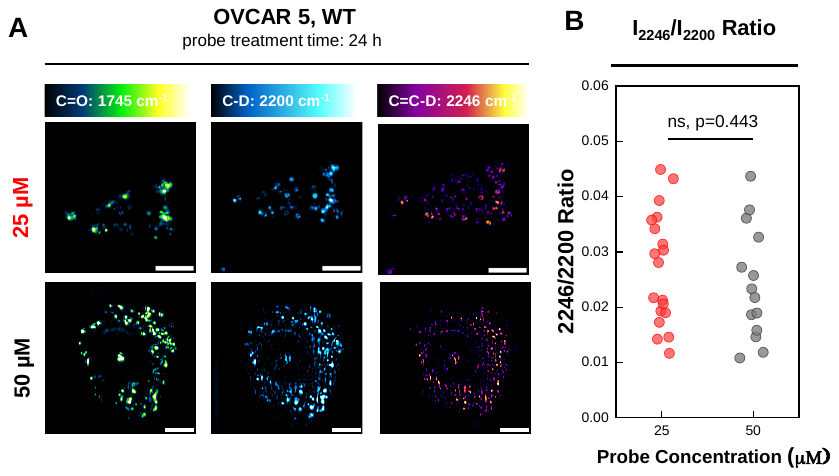

**Figure S9. PA-d31 concentration-dependent MIP analysis of the apparent SCD activity index.**

**(A)** Representative MIP images of OVCAR5 cells treated with 25 μM (top) or 50 μM (bottom) PA-d31 probe. Scale bar, 10 μm.

**(B)** Quantification of the 2246/2200 cm^-1^ intensity ratio in OVCAR5 cells treated with 25 μM (red) or 50 μM (black) PA-d31. 25: n = 19. 50: n =14, from N = 3 cells in each group, ns, p = 0.443.

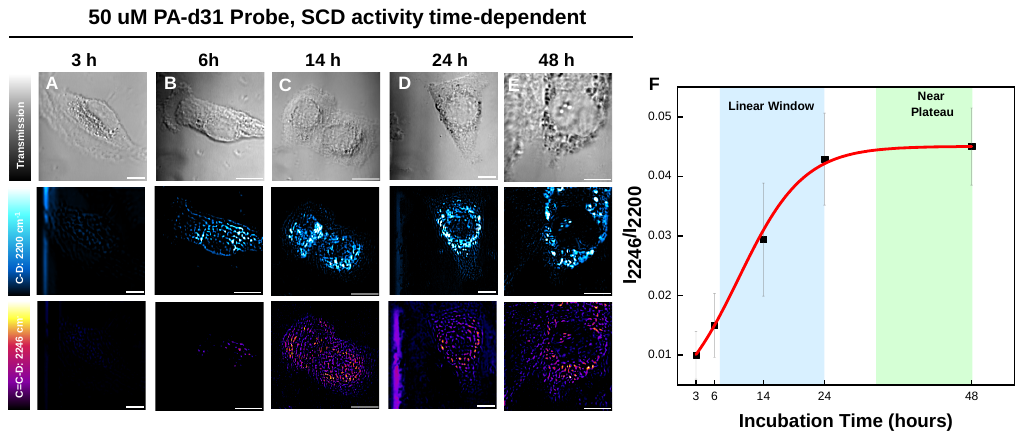

**Figure S10. Time-course analysis of intracellular SCD enzymatic activity.**

**(A)-(E).** Representative MIP images of cells for different incubation times at 2200 cm^-1^, 2246 cm^-1^ and corresponding transmission channel. Scale bar, 10 μm.

**(F)** Time -course analysis of the I_2246_/I_2200_ ratio at the time points shown in **(A)** to **(E)**. The red curve shows the fitted time-course curve with linear window and near-plateau window. N = 3 cells for each time point.

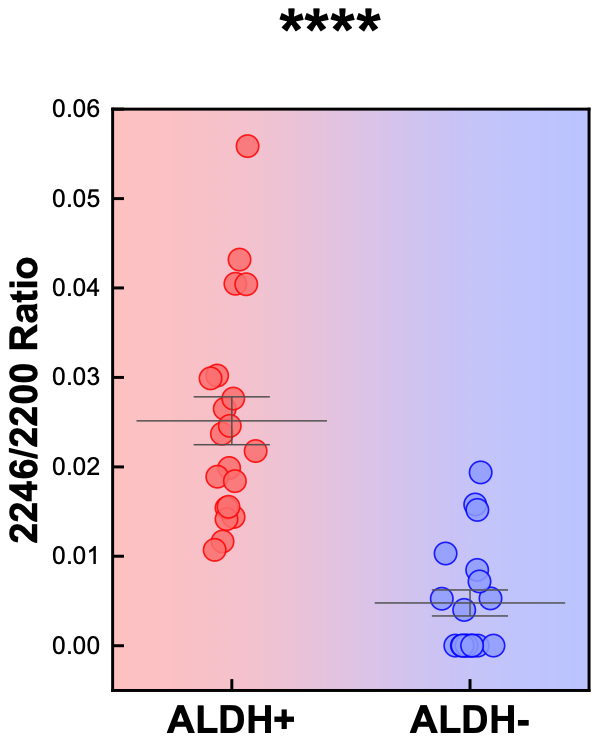

**Figure S11. Stem cell analysis of intracellular SCD activity index.** Comparison of the overall apparent SCD activity index (I_2246_/I_2200_) in ALDH+ and ALDH- groups. n = 19 in ALDH+ and n = 13 in ALDH- from N= 5 cells in each group, **** p =7.9×10^-5^.

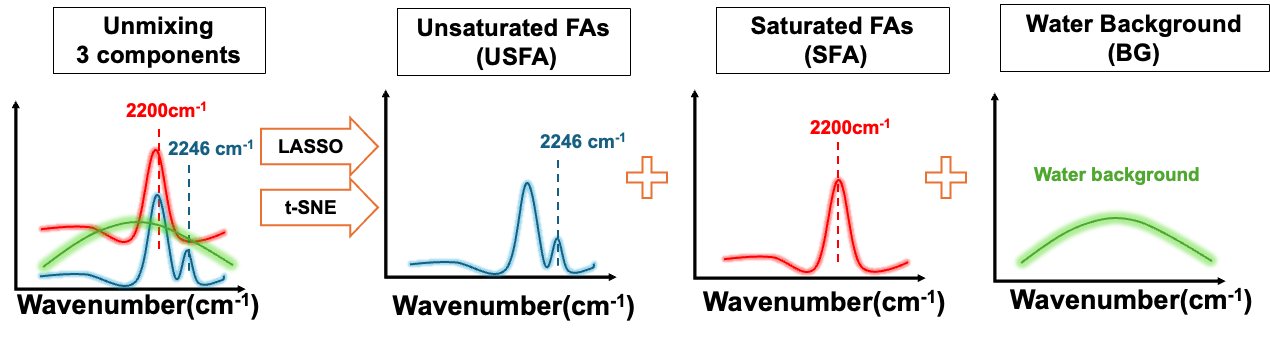

**Figure S12.** Schematic principle of LASSO unmixing method.

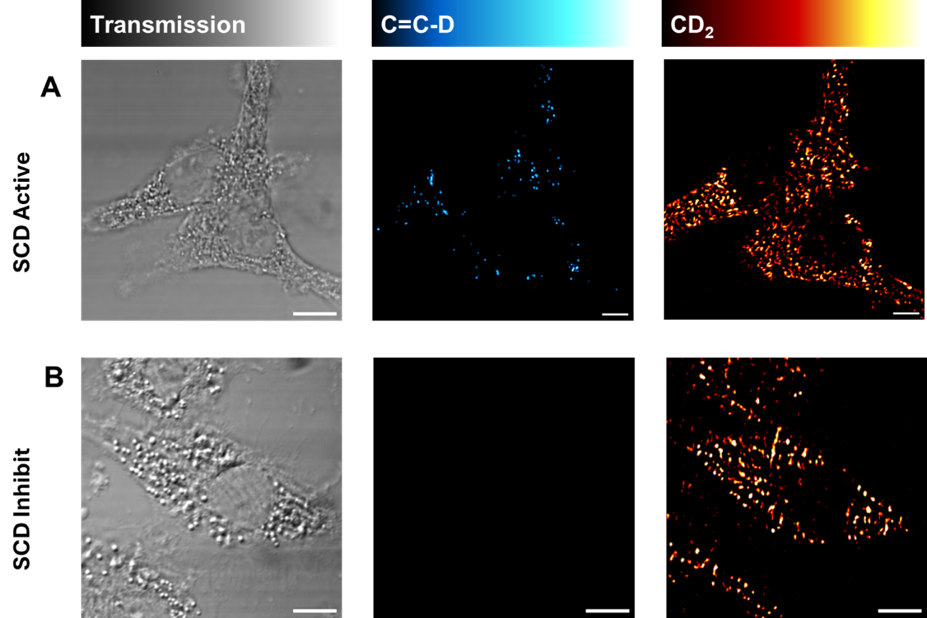

**Figure S13. LASSO unmixing result of (A)** SCD active group and **(B)** SCD inhibited cells. Scale bar, 10 μm.

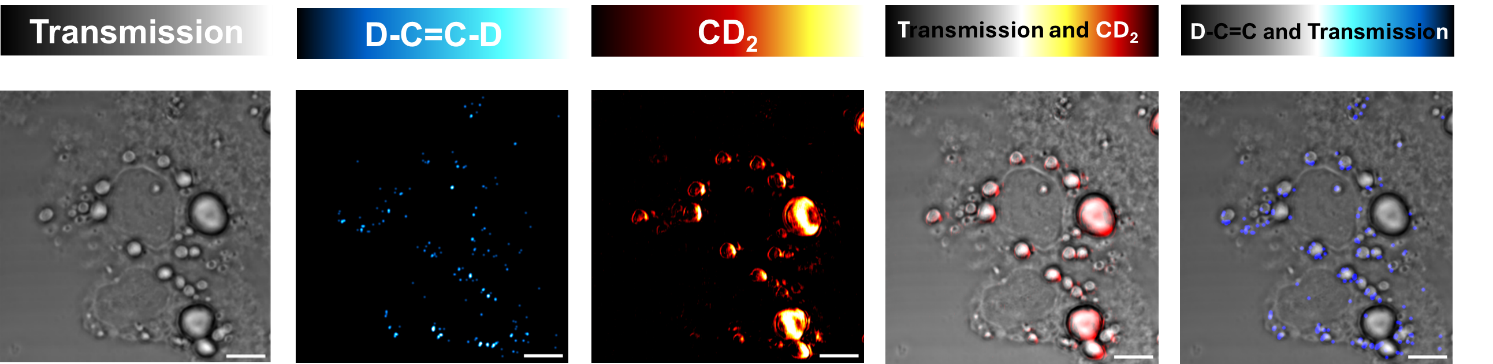

**Figure S14.** LASSO unmixing result of PA-d31 and OA co-treated OVCAR5 cells. Scale bar, 10 μm.

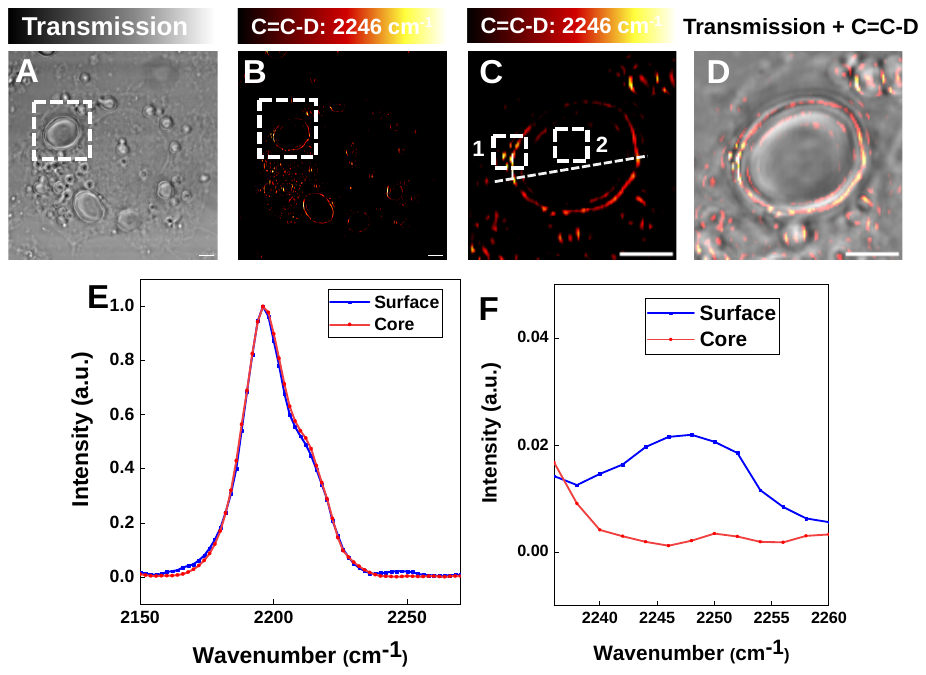
**Figure S15. MIP imaging of PA-d31 and OA co-treated OVCAR5 cells.**

**(A)** Transmission image of the treated cell.

**(B)** MIP image at 2246 cm⁻¹.

**(C)** Zoomed-in view of the dashed area in **(B)**.

**(D)** Merged image of the transmission and 2246 cm⁻¹ channels.

**(E)** MIP spectra from the LD surface region (ROI 1) and LD core region (ROI 2).

**(F)** Zoomed-in view of the 2246 cm⁻¹ region in **(E)**.

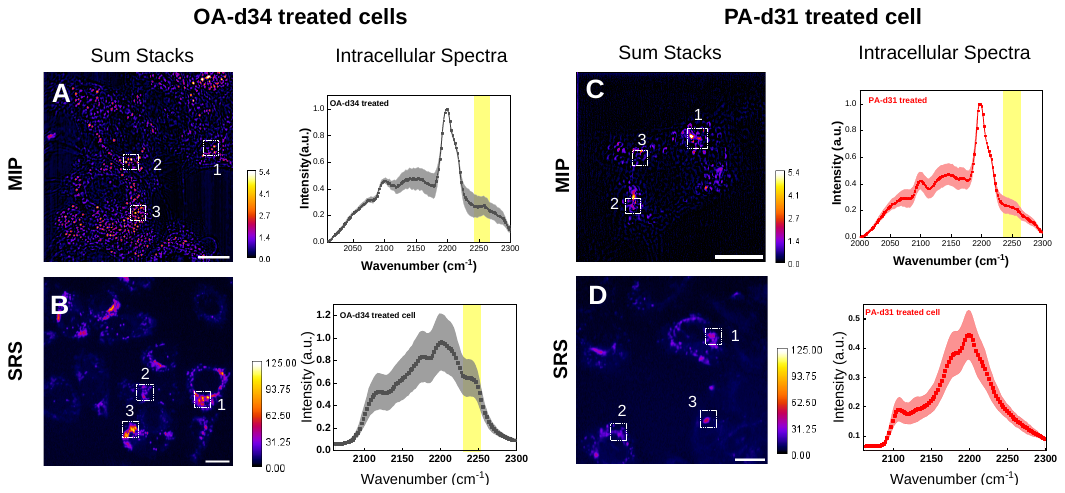

**Figure S16. MIP and SRS Hyperspectral Comparison on PA-d31 Treated SCD-active Cells.**

**(A)** and **(C)** MIP images Sum Stack (2000-2300 cm^-1^) of PA-d31 and OA-d34 treated SCD-active cell and its corresponding intracellular spectrum from selected ROIs.

**(B)** and **(D)** SRS images Sum Stack (2000-2300 cm^-1^) of PA-d31 and OA-d34 treated SCD-active cell and its corresponding intracellular spectrum from selected ROIs.
